# Modifications to a *LATE MERISTEM IDENTITY-1* gene are responsible for the major leaf shapes of cotton

**DOI:** 10.1101/062612

**Authors:** Ryan Andres, Viktoriya Coneva, Margaret H. Frank, John R. Tuttle, Sang-Won Han, Luis Fernando Samayoa, Baljinder Kaur, Linglong Zhu, Hui Fang, Daryl Bowman, Marcela Rojas-Pierce, Candace H. Haigler, Don C. Jones, James B. Holland, Daniel H. Chitwood, Vasu Kuraparthy

**Author notes:** These authors contributed equally to this work (Ryan Andres, Daniel H. Chitwood, Vasu Kuraparthy). Correspondence should be addressed to V.K.

## Abstract

Leaf shape is spectacularly diverse. As the primary source of photo-assimilate in major crops, understanding the evolutionary and environmentally induced changes in leaf morphology are critical to improving agricultural productivity. The role of leaf shape in cotton domestication is unique, as breeders have purposefully selected for entire and lobed leaf morphs resulting from a single locus, *okra* (*L-D_1_*). The *okra* locus is not only of agricultural importance in cotton (*Gossypium hirsutum* L.), but through pioneering chimeric and morphometric studies it has contributed to fundamental knowledge about leaf development. Here we show that the major leaf shapes of cotton at the *L-D_1_* locus are controlled by a HD-Zip transcription factor most similar to *Late Meristem Identity1 (LMI1)* gene. The classical *okra* leaf shape gene has133-bp tandem duplication in the promoter, correlated with elevated expression, while an 8-bp deletion in the third exon of the presumed wild-type *normal* leaf causes a frame-shifted and truncated coding sequence. Virus-induced gene silencing (VIGS) of this *LMI1-like* gene in an *okra* variety was sufficient to induce normal leaf formation. An intermediate leaf shape allele, *sub-okra*, lacks both the promoter duplication and the exonic deletion. Our results indicate that *sub-okra* is the ancestral leaf shape of tetraploid cotton and *normal* is a derived mutant allele that came to predominate and define the leaf shape of cultivated cotton.

## Introduction

Leaf shape is spectacularly diverse^1,2,3^. This diversity reflects evolutionary processes—either adaptive or neutral—that manifest through changes in developmental programming, environmental plasticity, or the interaction thereof^4^. As the primary sources of photo-assimilate in the world’s major crops, the role of leaves—their shapes, the constraints morphology places on other physiologically-relevant features, and the contributions of leaf shape to canopy and plant architecture—whether directly or indirectly selected upon during domestication, is an indisputably important consideration when discussing agricultural productivity. Although much is known about the developmental genetic basis of leaf morphology, only a handful of genes modifying leaf shapes in crops or responsible for natural variation among species have been identified^5-9^.

Cotton (*Gossypium* spp.) is the world’s most important source of natural fiber as well as a leading oilseed crop. The cultivated cottons (*G. hirsutum* and *G. barbadense*) are allotetraploid species (2n=4x=52, AADD) formed from the hybridization of diploids *G. arboreum* (2n=2x=26, AA) and *G. raimondii* (2n=2x=26, DD)^10^. Remarkable phenotypic diversity exists for leaf shape in cotton, widely ranging from entire (lacking dissection) to deeply lobed across both diploids and polyploids^11-13^. Leaf shape in *Gossypium* is an important agronomic trait that affects plant and canopy architecture, yield, stress tolerance, and other production attributes^14^. Among crops, leaf shape in cotton is unique; in recent history breeders used a single locus, *okra*, to purposefully alter leaf shape among cotton cultivars^14,15^. The four major leaf shapes of cotton: *normal*, *sub-okra*, *okra* and *super-okra* (**Fig. 1a**) are semi-dominant and allelomorphic at the L-D_1_ (*okra*) locus^14-20^ while *laciniate*, similar in morphology to *okra*, maps to the orthologous diploid A-genome locus (*L-A_1_*)^21^. Beyond agriculture, the *okra* locus is also of historical importance to leaf development. Not only was it used for one of the first comprehensive morphometric descriptions of leaves^11,22^, but pioneering studies creating *okra* chimeras determined the contributions of different cell layers to leaf shape^23,24^.

**Figure 1.**
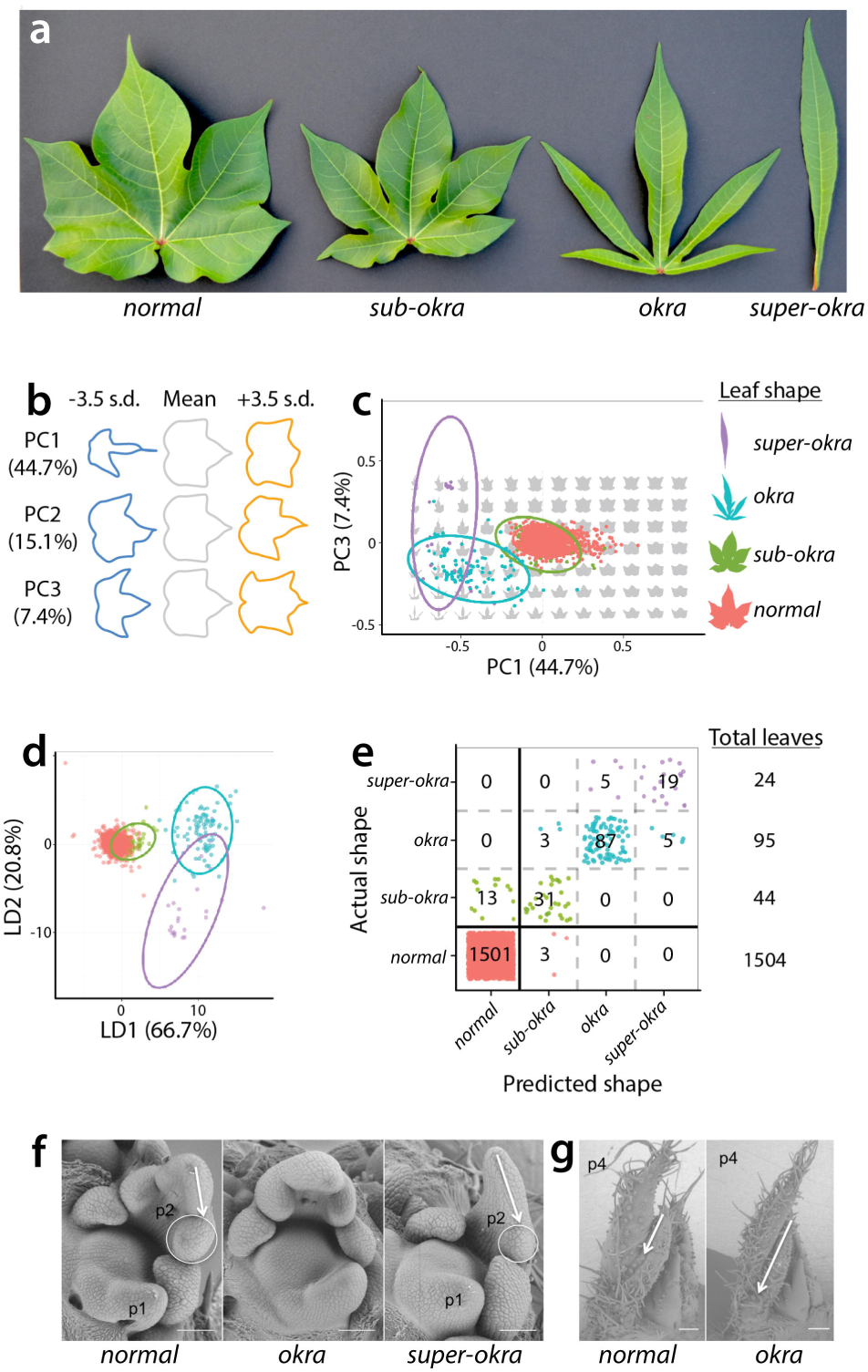
Morphometric analysis of leaf shapes conferred by the *sub-okra*, *okra*, and *super-okra* mutations. (**a**) Left to right, leaves representative of *normal*, *sub-okra*, *okra*, and *super-okra* leaf shape phenotypes. (**b**) Eigenleaves representative of leaf shapes found +/− 3.5 standard deviations along each Principal Component (PC) axis calculated from the harmonic series of an Elliptical Fourier Descriptor (EFD) analysis. Percent variance explained by each PC is provided. (**c**) PC1 and PC3 (PC2 not shown, as it explains asymmetric shape variance) separate *normal* and various *okra* leaf shape classes. 95% confidence ellipses provided. (**d**) Linear Discriminant Analysis (LDA) maximizes the discrimination of *normal* and *okra* leaf shape classes. LD1 and LD2 are shown, and the percent separation between phenotypic classes of leaves indicated. 95% confidence ellipses provided. (**e**) A confusion matrix, showing actual vs. predicted leaf shapes, constructed using linear discriminants. Leaf shape alone discriminants a majority of *normal* from *sub-okra*, *okra*, and *super-okra* leaf types. (**f**) Scanning Electron Micrographs (SEMs) of the Shoot Apical Meristem (SAM), P1, and P2 leaf primordia for *normal*, *okra*, and *super-okra* shoot apices. Note the displaced lobe in the P2 of *super-okra* relative to *normal*. (**g**) SEM of *normal* and *okra* shoot apices showing a more pronounced leaf lobe present by the P4 stage of leaf primordium development in *okra* relative to *normal*. Colors: *normal*, red; *sub-okra*, green; *okra*, blue; *super-okra*, purple.

The *okra* allelic series confers increasingly lobed leaf shapes from *sub-okra* to *okra* with the proximal lobes in mature *super-okra* reduced to a single linear blade (**Fig. 1a**). The characteristic shape of these four leaf morphs can be used both qualitatively and quantitatively^11,22^ to distinguish among their alleles (**Fig. 1b-e**). Classical development studies involving *okra* revealed the underlying factor acted early in leaf development in all tissue layers (L1, L2, L3) and cell autonomously^23,24^. Scanning Electron Micrographs (SEMs) of the shoot apical meristem show that the deeply lobed phenotype of *super-okra* is apparent by the P2 stage of leaf development (**Fig. 1f**), while the less severe *okra* manifests by the P4 stage (**Fig. 1g**).

The *L-D_1_* locus was placed on the short arm of chromosome 15-D_1_ (Chr15) using cytogenetics^25, 26^ and confirmed by QTL mapping^27-29^. The *L-D_1_* locus was localized to a 5.4cM interval near the telomere of Chr15^30^ and shuttle mapping utilizing the *laciniate* gene (*L-A_2_^L^*) from *G. arboreum* further reduced the candidate region to 112kb and ten genes^21^. Further, mapping and genomic targeting indicated two putative paralogous genes on Chr15 as the possible candidate genes ^30, 31^ for the *L-D_1_* locus. Here, we report the identification of a *LATE MERISTEM IDENTITY1 (GhLMI1-D1b)* gene, encoding an HD-Zip transcription factor, as the major determinant of leaf shape variation at the *L-D_1_* locus in cotton.

## Results

### The *okra* locus explains a majority of leaf shape variation in cultivated cotton

To determine the quantitative extent that the *okra* locus is responsible for controlling leaf shape in cultivated cotton, we morphometrically analyzed 1504 leaves from 420 cultivated cotton lines (**Supplementary Table 1**). The eigenleaves (representations of shape variance) resulting from a Principal Component Analysis (PCA) of the harmonic series of Elliptical Fourier Descriptors (EFDs) describe shape features associated with linear vs. palmatelylobed leaf types (PC1) and pronounced distal lobing (PC3), in addition to fluctuating asymmetry (PC2) (**Fig. 1b**). PC1 and PC3 (in addition to other PCs not shown) separate *normal* from *sub-okra, okra,* and *super-okra* leaf types and explain the majority of shape variance in the cotton accessions analyzed (**Fig. 1c**). A Linear Discriminant Analysis (LDA) performed to distinguish cotton accessions by leaf shape led to the following correct percentage of assignments to phenotypic class by EFDs alone: 99.8% *normal* (1501 of 1504 cases); 70.5% *sub-okra* (31 of 44 cases), 91.6% *okra* (87 of 95 cases) and 79.2% *super-okra* (19 of 24 cases) (**Fig. 1d, e**). The classification became 99% correct (1651 of 1667 cases) if only two classes were formed: *normal* and non-*normal* (inclusive of *sub-okra, okra, and super-okra*) (**Fig. 1e**).

Our results indicate that *okra* (*L-D_1_^O^*), a monogenic locus, is quantitatively responsible for the majority of leaf shape variance in cotton, the alleles of which can be discriminated from each other at high correct classification rates using shape information alone. A strongly monogenic basis for leaf shape in cotton is in contrast to a polygenic basis for leaf shape described in other crops^32,33^. Our results are consistent with classical morphometric work describing the profound role the *L-D_1_^O^* locus plays in determining cotton leaf shape^11,22^.

### Fine mapping of the *L-D_1_* locus in a large F_2_ population

1,027 F_2_ plants from the cross NC05AZ21 × NC11-2100 showed the expected phenotypic ratio for single gene inheritance of *okra* leaf shape (**Supplementary Table 2)**. Genotyping with the co-dominant flanking SSRs of *L-D_1_* (Gh565 and DPL0402) identified 122 recombinants and produced a genetic map (**Fig. 2a**) similar to the preliminary mapping^30^. Genotyping of these recombinants showed marker 13-LS-195 continued to co-segregate with leaf shape phenotype, confirming that 13-LS-195 is tightly linked to the leaf shape locus (*L-D_1_*) in cotton^30^. The resulting 337 kb, 34 gene candidate interval (**Fig. 2b**)^30^ was further resolved to 112 kb and ten genes using orthologous mapping of the homeologous *laciniate* gene^21^ (**Fig. 2c**).

**Figure 2.**
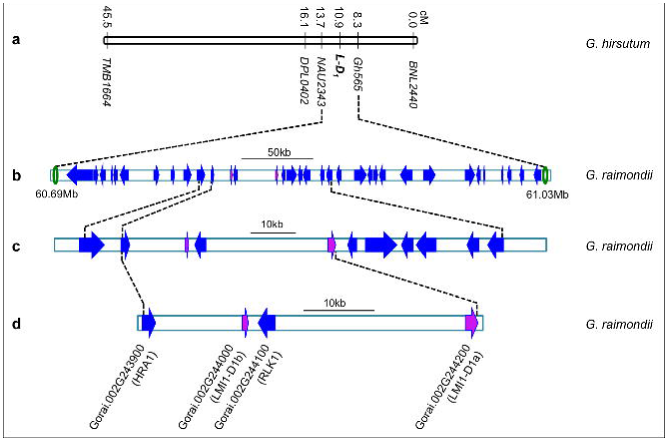
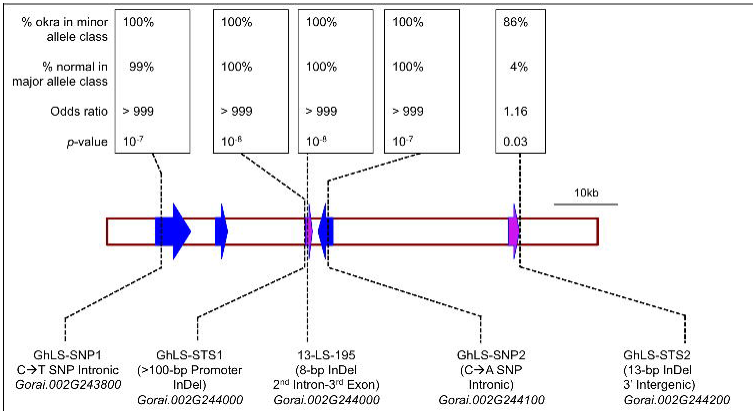
Genetically resolving the *L-D_1_* locus in Upland cotton. (**a**) Genetic mapping of *L-D_1_* locus based on bi-parental mapping. (**b**) Orthologous mapping to the sequenced D genome donor (*G. raimondii*) chromosome 2 (337kb, 34 putative genes)^21^. (**c**) Shuttle mapping utilizing the orthologous *laciniate* (*L-A_2_*) locus from donor diploid *G. arboreum* (112kb, 10 putative genes)^12^. (**d**) Fine mapping combining an association mapping panel and two sets (BC_8_ and BC_3_) of isogenic lines (52kb, putative 4 genes). (**e**) Association analysis statistics, adjusting for population structure for variants within candidate gene region.

### Fine mapping using association mapping and isogenic lines

#### Novel markers

Using *G. raimondii* sequence information^34^, four novel markers were developed within the ten gene candidate region (**Supplementary Fig. 1**): 1) a large (>100 bp) STS marker (GhLS-STS1) in the promoter region of *Gorai.002G244000*, 2) an 8-bp STS marker (13-LS-195) within *Gorai.* 002G244000, 3) a C→A SNP (GhLS-SNP2) in the intron of *Gorai.002G244100* and 4) a 13-bp STS marker (GhLS-STS2) <300 bp downstream of the *Gorai.002G244200* stop codon. An additional SNP marker, (GhLS-SNP1) in the intron of *Gorai.002G243800* was also designed despite the fact that this gene had already been excluded by orthologous mapping^21^ (**Fig. 2c**).

#### Association mapping

Of the four markers used to genotype the diversity panel of 538 accessions, three markers (GhLS-STS1, 13-LS-195, and GhLS-SNP2) showed complete association with leaf shape while marker GhLS-STS2 showed no association between the two most common leaf shapes, *normal* and *okra* (**Supplementary Table 3**). The lack of association of this STS marker with leaf shape was sufficient to reduce the ten gene, 112-kb candidate region (**Fig. 2c**) to a four gene, 52 kb region between *Gorai.002G243900* and *Gorai.002G244200* (**Fig. 2d** and **Supplementary Table 3**).

Population structure was estimated in a subset of 404 lines of the 544 member panel using SSR markers distributed throughout the genome. Association tests of variants in the candidate gene region and of SSRs throughout the genome confirmed that the four remaining candidate genes showed very strong and significant (*p* < 10^-7^) association with leaf shape after correcting for population structure (**Fig. 2e** and **Supplementary Table 4**). A few SSR markers also had significant associations, which may have occurred by chance when rare allele classes were observed in okra types (**Supplementary Fig. 2a** and **Supplementary Table 4)**. After fitting the most significant candidate gene marker as a covariate and re-testing the background markers for associations, no markers outside the candidate gene interval were significant (**Supplementary Fig. 2b** and **Supplementary Table 4**).

#### Mapping using isogenic lines

We confirmed that genes conferring the *okra* phenotype in the mapping parent NC05AZ21 and *okra* isogenic line (LA213-*okra*) are allelic by phenotyping their F_1_ (**Supplementary Fig. 3**) and an F_2_ population. Genotyping of two sets of isolines (BC_8_ in Stoneville 213 and BC_3_ in Stoneville 7A backgrounds) with the four novel markers showed a similar marker pattern as observed in the association mapping (**Supplementary Table 3** and **Supplementary Table 5**), confirming the resolution of the candidate region to four genes and 52 kb.

**Figure 3.**
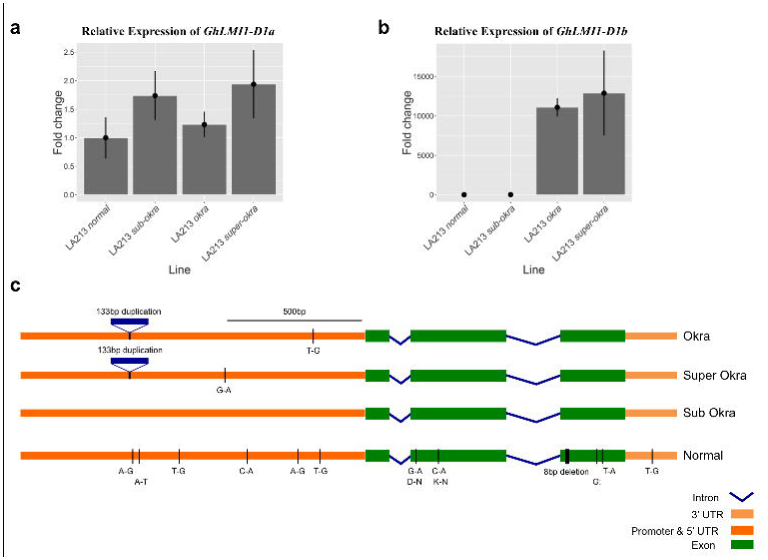
Nucleotide polymorphisms of *GhLMI1-D1b* gene and expression analysis of candidate genes using qRT-PCR among different leaf shapes. (**a, b**) Relative transcript levels of leaf shape candidate genes (*GhLMI1-D1* a and *GhLMI1-D1b*) among isolines at ∼90 days after planting. There is no difference in the expression of *GhLMI1-D1a* among leaf shapes but a significant increase of *GhLMI1-D1b* expression in *okra* and *super-okra*. Error bars represent the standard deviation of the fold change. (**c**) Polymorphisms among *GhLMI1-D1b* sequences from the four major leaf shapes. The 133bp tandem promoter duplication was found only in *okra* and *super-okra* and in parallel with elevated gene expression. The 8bp deletion was found only in *normal* and causes a frameshift mutation and premature stop codon. All other polymorphisms are SNPs with unknown effect on the gene expression and protein function. *Sub-okra* is set as the standard to which the other three leaf shapes are compared.

Thus, characterizing the association-mapping panel and isolines with novel markers reduced the candidate genomic region to 52kb containing four genes. Of the four genes, *Gorai.002G244200* (hereafter *GhLMI1-D1a*) and *Gorai.002G244000* (hereafter *GhLMI1-D1b*) are paralogues coding for HD-Zip transcription factors with 71.2% protein similarity. Their homologs were implicated in flowering time and leaf complexity in Arabidopsis^35,8,9^. Of the remaining two putative genes, *Gorai002G244100* (hereafter *GhRLK1*), is a serine/threonine protein kinase while *Gorai.002G243900 (* hereafter *GhHRA1)* is a trihelix transcription factor.

### Expression analysis of leaf shape gene candidates

Expression analysis was performed to help identify the causal gene among the four remaining candidates. Semi-quantitative expression analysis revealed that neither *GhHRA1* nor *GhRLK1* were expressed in young leaf tissue (**Supplementary Fig. 4**). Based on this observation together with their lack of sequence polymorphisms from novel STS marker development, we eliminated these two genes from consideration. *GhLMI1-D1a* was similarly expressed among leaf shapes while *GhLMI1-D1b* expression was detected only in *okra* and *super-okra* (**Supplementary Fig. 4**). qRT-PCR confirmed the equivalent expression of *GhLMI1-D1a* among leaf shapes while *GhLMI1-D1b* was upregulated in *okra* and *super-okra* compared to *normal* and *sub-okra* (**Fig. 3a, b**). This indicated that up-regulation and/or ectopic expression of *GhLMI1-D1b* in *okra* and *super-okra* may be responsible for these leaf shapes.

### Sequencing of *GhLMI1*-like genes

To identify DNA polymorphisms among leaf shapes, *GhLMI1-D1a* and *GhLMI1-D1b* were sequenced in 20 cotton varieties (**Supplementary Table 6**) and comparisons were made relative to *sub-okra* (**Fig. 3c** and **Supplementary Fig. 5**). Sequence analysis of *GhLMI1-D1a* identified two variants (**Supplementary Fig. 5)**. Variant 1 was found in two *normal* and all five *okra* and *sub-okra* lines. Variant 2 was found in the remaining three *normal* and all five *super-okra* lines. The 13-bp InDel (GhLS-STS2), which showed no association with leaf shape in the association analysis (**Supplementary Table 3**), is placed ∼100 bp from the end of the 3’ UTR. Although this polymorphism lies outside *GhLMI1-D1a*, the proximity, lack of major polymorphisms, and identical expression among leaf shapes provide evidence against *GhLMI1-D1a* as the candidate gene underlying *L-D_1_*.

Sequence analysis of *GhLMI1-D1b* identified two prominent polymorphisms among leaf shapes. First, a 133-bp tandem duplication located ∼800 bp upstream of the transcription start site was unique to *okra* and *super-okra* (**Figure 3c**) and may explain the altered expression of *GhLMI1-D1b* seen earlier (**Fig. 3b**). The second notable polymorphism was an 8-bp deletion in the 3^rd^ exon of *GhLMI1-D1b* found only in *normal* (**Fig. 3c**). The exonic location of this deletion was confirmed through Sanger sequencing of *GhLMI1-D1b okra* cDNA. Translation of the resulting *normal* coding sequence produces a frameshift 156 amino acids (aa) into the protein and a premature stop codon truncating the protein from 228 aa to 178 aa (**Fig. 5a**). Neither of the conserved functional domains of an HD-Zip transcription factor appear directly impacted by the deletion. However, the frameshift introduces multiple leucines that may disturb the characteristic spacing of the leucine zipper and alter protein-protein interactions (**Fig. 5a**).

**Figure 4.**
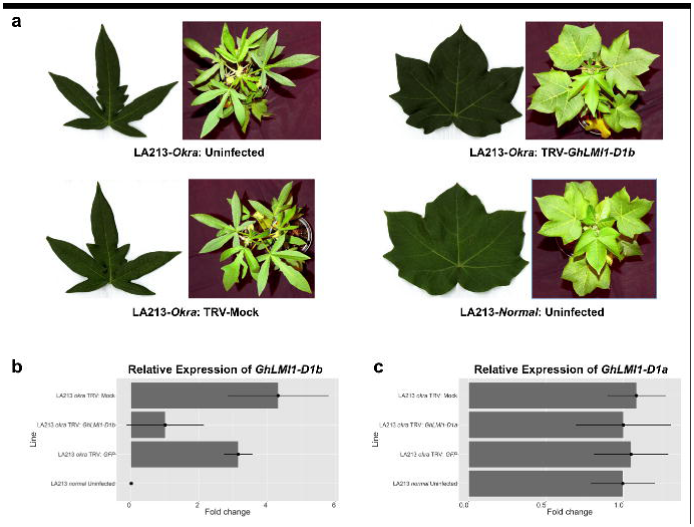
Functional characterization of *GhLMI1-D1b* using virus-induced gene silencing (VIGS). (**a**) Representative 6^th^ true leaf and 4 week old plants from VIGS experiment showing the reversion to *normal* leaf shape in the *GhLMI1-D1b* silencing treatment. (**b**) Relative transcript levels of candidate genes in the *GhLMI1-D1b* silenced and control LA213-*Okra* plants confirmed the effective knockdown of *GhLMI1-D1b* **. (c)** Transcript levels of *GhLMI1-D1a* were unaffected by VIGS treatment, confirming silencing was specific to *GhLMI1-D1b*.

**Figure 5.**
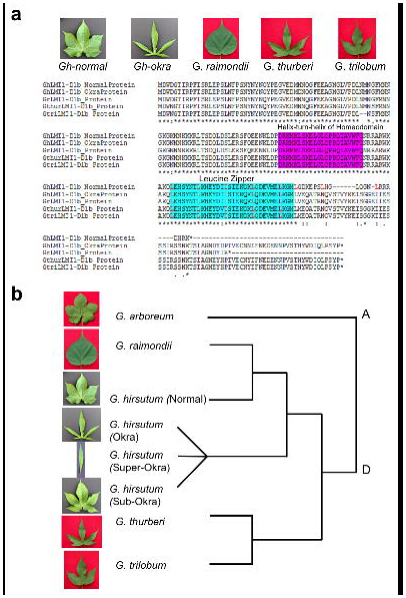
Functional prediction and phylogenetic analysis of *LMI1*-like genes among different cotton leaf shapes. (**a**) Amino acid translations in the *Gossypium LMI1-D1b* alleles. Normal in *G. hirsutum* and simple leaf in *G. raimondii* show truncated proteins while *G. hirsutum-okra*, *G. trilobum* and *G. thurbeii* with dissected leaves code for functional *LMI1-D1b* protein. Frameshift mutation resulting from 8bp deletion in *normal* introduces additional leucines (L) at 7 amino acid intervals (highlighted in red) that may interfere with the functionality of this domain. (**b**) Phylogenetic analysis showing the close relationship between *okra*, *sub-okra*, and *super-okra* but not to *G. thurberi* or *G*. *trilobum*. Conversely, *normal* CDS appears more closely related *G. raimondii*.

Both major polymorphisms (GhLS-STS1 for promoter duplication and 13-LS-195 for 3^rd^ exon InDel) were completely associated with leaf shape in the association panel and isogenic lines (**Supplementary Table 3** and **5**). Taken together with the gene expression differences, these results provided strong evidence that modifications to *GhLMI1-D1b* are responsible for the various leaf shapes of cotton.

### Functional characterization of *GhLMI1-D1b* using virus-induced gene silencing (VIGS)

As expression analysis indicated the elevated or ectopic expression of *GhLMI1-D1b* could be responsible for okra leaf shape (**Fig. 3b**), we hypothesized that silencing of *GhLMI1-D1b* in *okra* would reduce transcript levels and confer a simpler leaf shape. A 461bp fragment of *GhLMI1-D1b^Okra^* encompassing 268bp of the coding sequence (CDS) and 193bp of the 3’ untranslated region (UTR) was used in VIGS. Including the 3’ UTR sequence was expected to minimize off-target silencing of other *GhLMI1-* like genes, especially *GhLMI1-D1a.* Silencing of *GhLMI1-D1b* in an *okra* isoline led to a pronounced reduction in leaf lobing compared with uninfected and negative controls and a brief period of normal leaf production (**Fig. 4a** and **Supplementary Fig. 6**). Silencing of *GhLMI1-D1b* was eventually overcome, leading to a reversion to *okra* in a timeframe similar to the loss of silencing seen in the TRV: *CHLI* positive control (**Supplementary Fig. 6**).

The level of *GhLMI1-D1b* transcript was substantially reduced in the TRV: *GhLMI1-D1b* treatment compared to the negative controls (**Fig. 4b)**. This proved that knocking down the *GhLMI1-D1b* transcript through VIGS was sufficient to induce *normal* leaf formation in an *okra* variety. Furthermore, expression of *GhLMI1-D1a* was unaffected by VIGS (**Fig. 4c**) demonstrating specificity to *GhLMI1-D1b*. Thus, phenotyping and transcript profiling of TRV: *GhLMI1-D1b* leaves confirmed that altered expression of *GhLMI1-D1b* was responsible for the *okra* leaf shape of cotton.

### Phylogenetic analysis of *LMI1*-like genes in diploid and tetraploid cotton

Outside of the promoter duplication *sub-okra*, *okra*, and *super-okra GhLMI1-D1b* were identical except for one promoter SNP unique to both *okra* and *super-okra* (**Fig. 3c**). Phylogenetic analysis confirmed the close sequence relatedness of the *sub-okra*, *okra*, and *super-okra GhLMI1-D1b* alleles (**Fig. 5b**). However, *normal GhLMI1-D1b* was considerably different from the other alleles with six unique promoter SNPs and two SNPs in the second exon, both of which cause amino acid changes. *Normal* also carries a 1 bp deletion in the 3^rd^ exon, an additional third exon SNP, and a SNP in the 3’ region (**Fig. 3c**).

The 1-bp deletion would also cause a frameshift and premature stop codon if not for the preceding 8-bp deletion. Interestingly, this 1-bp deletion is identical to one carried by the D genome donor *G. raimondii* in *Gorai.002G244000.* While it is unclear if the 1-bp deletion in *Gorai.002G244000* plays any role in the entire leaf shape of *G. raimondii* (**Fig. 5a**), it would be interesting to determine the leaf shape of a tetraploid carrying just the 1-bp deletion without the preceding 8-bp deletion *. Normal* leaf *GhLMI1-D1b* also shares many of its other polymorphisms with *Gorai.002G244000* including four out of the six promoter SNPs, the G→A SNP in the 2^nd^ exon, the T A SNP in the third exon, and the 3’ UTR SNP. Phylogenetic analysis shows that *normal* leaf *GhLMI1-D1b* CDS is more closely related to the *G. raimondii LMI1-D1b* gene than to the alleles in other tetraploids with variable leaf shapes (**Fig. 5b**). *GaLMI1-A1b* from moderately lobed *G. aboreum* is predicted to produce a full-length *LMI1-*like protein that is identical in length to those produced by the non-*normal* leaf shapes of tetraploid cotton.

In order to further assess the variability of *LMI1-D1b* genes in *Gossypium*, *GhLMI1-D1b* was Sanger sequenced in two additional D genome diploids, *G. thurberi* and *G. trilobum*. Both of these species have highly lobed leaves similar to *G. hirsutum okra* (**Fig. 5a**). Both of the *G. thurberi* and *G. trilobum LMI1-D1b* genes were full-length and similar to those found in the *G. hirsutum* leaf shape mutants and *G. arboreum*. Thus, while only a handful of species from the large *Gossypium* genus have been analyzed, there exists a trend that species with lobed leaves carry full-length *LMI1*-*D1b* genes while those with entire leaves produce truncated *LMI1-1b* genes (**Fig. 5a**). Extending beyond *G. hirsutum*, this provides evidence for a broad role of *LMI1-1b* genes in controlling the diversity of leaf shapes seen throughout *Gossypium.*

### Transcriptome-wide changes accompanying *GhLMI1-D1b*

The morphological effects of *okra* leaf shape are evident by the P4 leaf primordia stage (**Fig. 1g)**. To analyze gene expression changes preceding observable changes in leaf morphology, we used RNA-Seq to determine transcriptional changes at the P2 stage of leaf development (**Fig. 6a**). 453 genes are differentially expressed between *okra* and *normal* BC_8_ isolines (**Fig. 6b** and **Supplementary Table 7**). Gene Ontology (GO) enrichment analysis for genes down regulated in *okra* indicated an association with microtubule and kinesin processes, suggesting a cellular basis for the observed changes in leaf morphology (**Supplementary Table 8**). Most of the GO terms for genes up regulated in *okra* relative to *normal* are related to photosynthesis (**Supplementary Table 9**). Previous transcriptomic analyses in tomato apices revealed photosynthetic-related gene expression and genetic changes in leaf complexity are positively correlated^33^. The direction of this correlation is the same in *okra* (i.e., positively correlated with more dissected leaf morphology) implying possible connections between leaf development and the capacity for photosynthesis.

**Figure 6.**
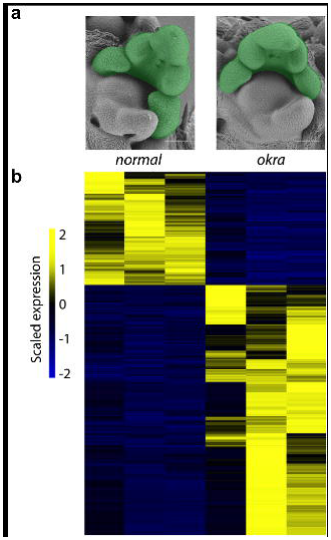
Transcriptomic comparison of *okra* versus *normal* P2 stage primordia. (**a)** SEMs of *normal* and *okra* shoot apices. P2 stage leaf primordia (highlighted in green) were hand dissected and processed for Illumina RNA-sequencing. **b)** Heatmap visualization of the 453 genes that are significantly differentially expressed (false discovery rate ≤ 0.05) between *normal* and *okra* P2 samples (**Supplementary Table 7-9**). Scaled and centered reads per million values for *normal* and *okra* biological replicates are plotted in the left three columns and right three columns, respectively. Yellow indicates up regulated and blue indicates down regulated expression values. Scale bars = 100 micrometers.

### Subcellular/Nuclear localization of *GhLMI1-D1b*

Transient expression of *35S::GhLMI1-D1b^Okra^-GFP* and *pUBQ10::GhLMI1-D1b^Okra^-GFP* in *Nicotiana benthamiana* was used to determine the subcellular localization of *GhLMI1-D1b^Okra^. GhLMI1-D1b^Okra^-GFP* was detected in the nucleus of transformed tobacco plants (**Supplementary Fig. 7**) in a pattern that co-localized with the nuclear marker RFP-Histone2B (**Supplementary Fig. 7**). The nuclear localization of *GhLMI1-D1b^Okra^-GFP* is consistent with the predicted function of *LMI1*-like genes as transcription factors^35,8,9^.

## Discussion

Leaf shape varies dramatically across plant evolution and in response to the environment. Understanding the genetic architecture of leaf shape is critical for harnessing its variation to modify plant physiology and improve agronomic profitability^2,4^. Using a diverse array of genomic and molecular tools, we established that the multi-allelic, major leaf shape locus *L-D_1_* of cotton is governed by *GhLMI1-D1b* encoding an HD-Zip transcription factor.

*LMI1*-like genes, in particular their duplication and regulatory diversification, have recently been proposed as evolutionary hotspots for leaf shape diversity in model plants^8,35^. The evidence presented here indicates that the major leaf shapes of cultivated cotton are controlled by the same pathway. The near-tandem duplication of *LMI1-*like genes in *Gossypium* (**Fig. 2d**) are unique from those previously described, as the Malvales and Brassicales diverged prior to duplications observed in the Brassicaceae^8,9^. Therefore, the separate *LMI1*-like duplication event in *Gossypium* indicates convergent evolution and strengthens the evidence that *LMI1*-like genes are evolutionary hot spots for modifying leaf shape^8,9^.

Sequence analyses of *GhLMI1-D1b* in a set of twenty cultivars elucidated two major polymorphisms among different leaf shapes at the *L-D_1_* locus (**Fig. 3c** and **Supplementary Table 5, 6**). First, an 8-bp deletion near the beginning of the third exon was found only in *normal GhLMI1-D1b*. This deletion results in a frameshift and premature stop codon in the predicted *normal GhLMI1-D1b* protein that may interfere with the function of the leucine zipper motif (**Fig. 5a**). Additionally, the C-terminal truncation may impact protein stability by removing residues necessary for proper folding. The second co-segregating polymorphism was a 133bp tandem duplication ∼800bp upstream of the *GhLMI1-D1b* transcription start site (**Fig. 3c** and **Supplementary Table 5**). This duplication was present in all *super-okra* and *okra* alleles but absent from all *normal* and *sub-okra* alleles. Furthermore, expression of *GhLMI1-D1b* was upregulated in plants with *okra* and *super-okra* leaf shapes (**Fig. 3b**). This finding is consistent with previous reports that promoter modifications to *LMI1-*like genes alter expression and leaf shape in model plants^8,9^. Finally, the down-regulation of *GhLMI1-D1b* by VIGS in a cotton accession with the *okra* leaf phenotype strongly reduced leaf lobing and resulted in the production of *normal* leaves (**Fig. 4a).**

Based on phylogenetic analysis, it appears that *normal* originated in *G. raimondii LMI1-D1b* (**Fig. 5b**). Owing to their shared 1-bp deletion, frameshift, and premature stop codon, this protein may already have been non- or partially functional. The subsequent 8-bp deletion, which causes an earlier frameshift and premature stop codon, would be expected to even further compromise the protein function. Sequence analysis indicated the other three leaf shape alleles are not derivatives of *G. raimondii LMI1-D1b.* This ancestral allele may have derived from a lobed D genome diploid, although likely not *G. thurberi* or *G. trilobum* (**Fig. 5b**). Once the *sub-okra* allele developed within the D genome, it likely gave rise to *okra* via a single duplication of 133bp in the promoter region. The origin of this promoter duplication is currently unclear but it may have arisen via unequal crossing over or gene conversion. Consistent with the expression results and gene silencing results obtained here, this promoter duplication led to ectopic and/or over-expression of *GhLMI1-D1b* and an increase in the degree of leaf lobing and complexity. The finding of only a single additional promoter SNP in the entire *GhLMI1-D1b* genic region between *sub-okra* and *okra* indicates this event happened relatively recently. Only two promoter SNPs differentiate *super-okra* from *okra* (**Fig. 3c**) consistent with the knowledge that *super-okra* originated from *okra* within the last 90 years^18^. Phylogenetic analysis involving sequences from all *Gossypium* species would establish the evolution and adaptive significance of *LMI1* genes in cotton.

*Okra* is an exceptional mutation affecting leaf development. It inspired an early quantitative framework for leaf shape across evolution and development^11,22^ and through chimeric studies, revealed some of the first insights into the morphogenetic and cellular basis of leaf morphology^243,24^. In two separate instances within the Brassicaceae, the homologs of *GhLMI1-D1b* have been implicated in evolutionary shifts in leaf morphology ^8,9^. That the *okra* locus confers a monogenic basis to most of the leaf shape variance in cotton, the mechanisms for which have been studied in other model organisms, and that it is implicated in the productivity of a major crop, demonstrates a unique intersection between agriculture and the evolutionary and developmental basis of leaf morphology.

## URLs

Clustal Omega Multiple Sequence Alignment, http://www.ebi.ac.uk/Tools/msa/; *G. raimondii* genome, www.phytozome.net; mRNA Translation to Protein Sequence, http://web.expasy.org/translate/; FancyGene, http://bio.ieo.eu/fancygene/; Helix-turn-helix Prediction, https://npsa-prabi.ibcp.fr/cgi-bin/primanal_hth.pl; Leucine Zipper Prediction, http://2zip.molgen.mpg.de/; Cotton Database, www.cottongen.org; https://gtac.wustl.edu/; Sequence Alignment, http://samtools.sourceforge.net; Read mapping, http://bedtools.readthedocs.org/en/latest/content/tools/multicov.html; Pairwise gene comparison, https://bioconductor.org/packages/3.0/bioc/html/edgeR.html.

## METHODS

Methods and any associated references are available in the online version of the paper.

*Note: Supplementary information is available in the online version of the paper.*

## AKNOWLEDGEMENTS

Cotton Incorporated supported this work through its core-funding and PhD Fellowship program. For additional support of this research, we thank the NC Cotton Growers Association Inc. and the National Science Foundation: grants IOS-1238014 and IOS-1025947 supported the work of LFS and JRT, respectively. We appreciate the technical help by Jared Smith and Sharon Williamson USDA-ARS and Cathy Herring of NCDA. We appreciate Drs. Richard Percy and James Frelichowski of the USDA-NCGC for supplying the cotton germplasm.

## AUTHOR CONTRIBUTIONS

V.K., R.A. and D.C. designed the research and wrote the manuscript, with contributions and final review by the other authors. R.A. and V.K. performed fine mapping, gene cloning and functional genomic analyses. V.C., M.F. and D.C. performed morphometric, SEM and RNA-Seq analyses. L.F.S. and J.B.H. performed association analyses. L.Z., B.K., H.F. genotyped the diversity panel. R.A., J.R.T., C.H.H. and V.K. performed VIGS experiments. M.R.P. and S.W.H. performed the tobacco transient expression assay.

## Supplementary figures

**Supplementary Figure 1.**
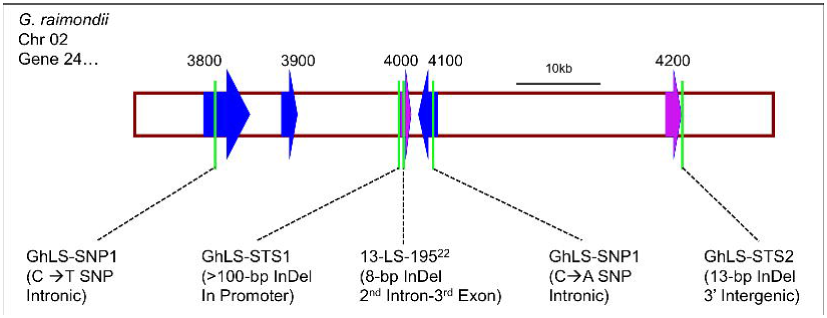
Markers used in association mapping and isogenic line studies. Four novel markers: GhLS-STS1, GhLS-STS2, GhLS-SNP1 and GhLS SNP2 were developed and run in both studies along with 13-LS-195 that had previously co-segregated with leaf shape phenotype^21^. Four markers: GhLS-STS1, 13-LS-195, GhLS-SNP1 and GhLS-SNP2 showed complete association with leaf shape phenotype. GhLS-STS2 showed no association with leaf shape.

**Supplementary Figure 2.**
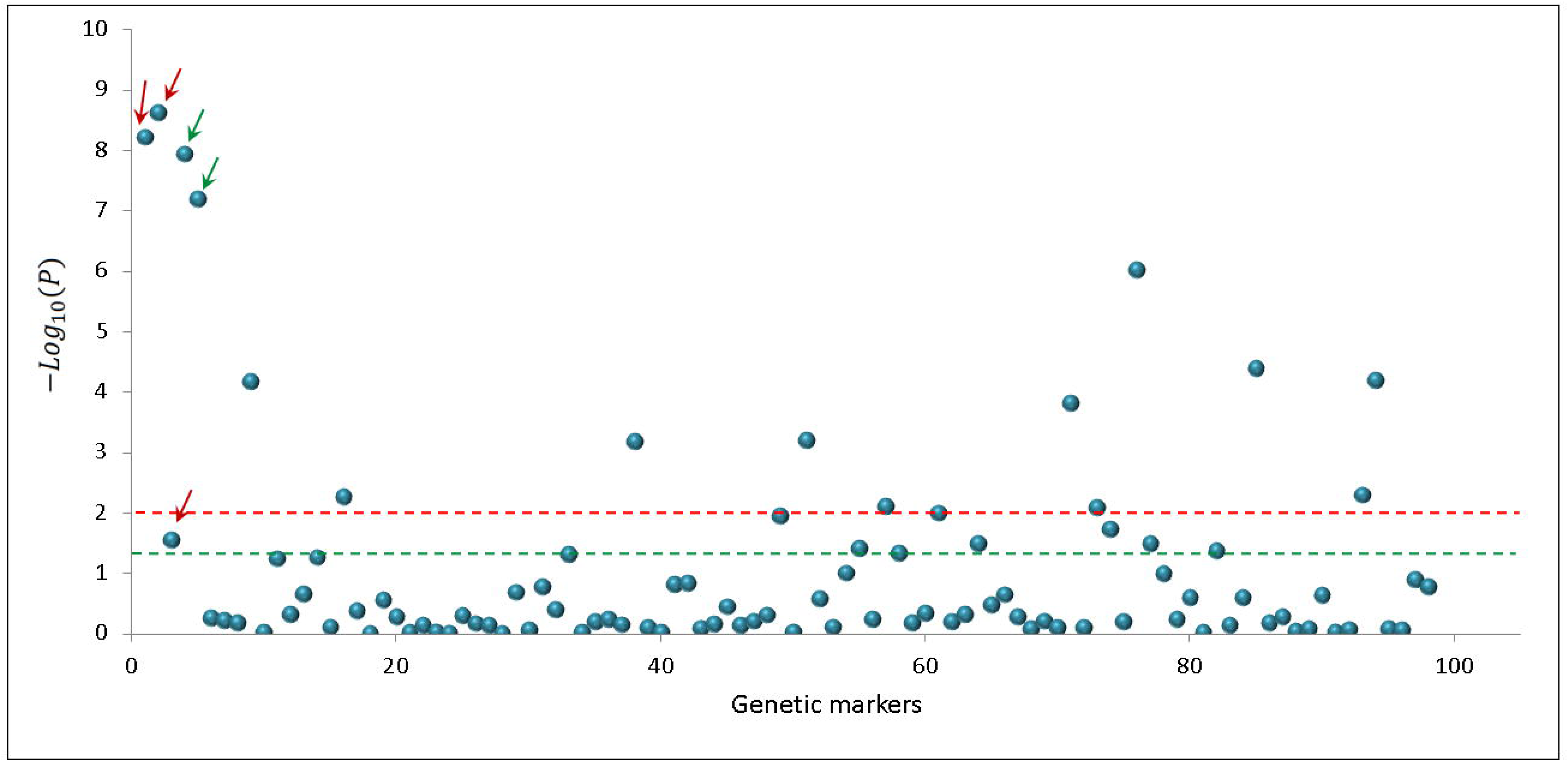
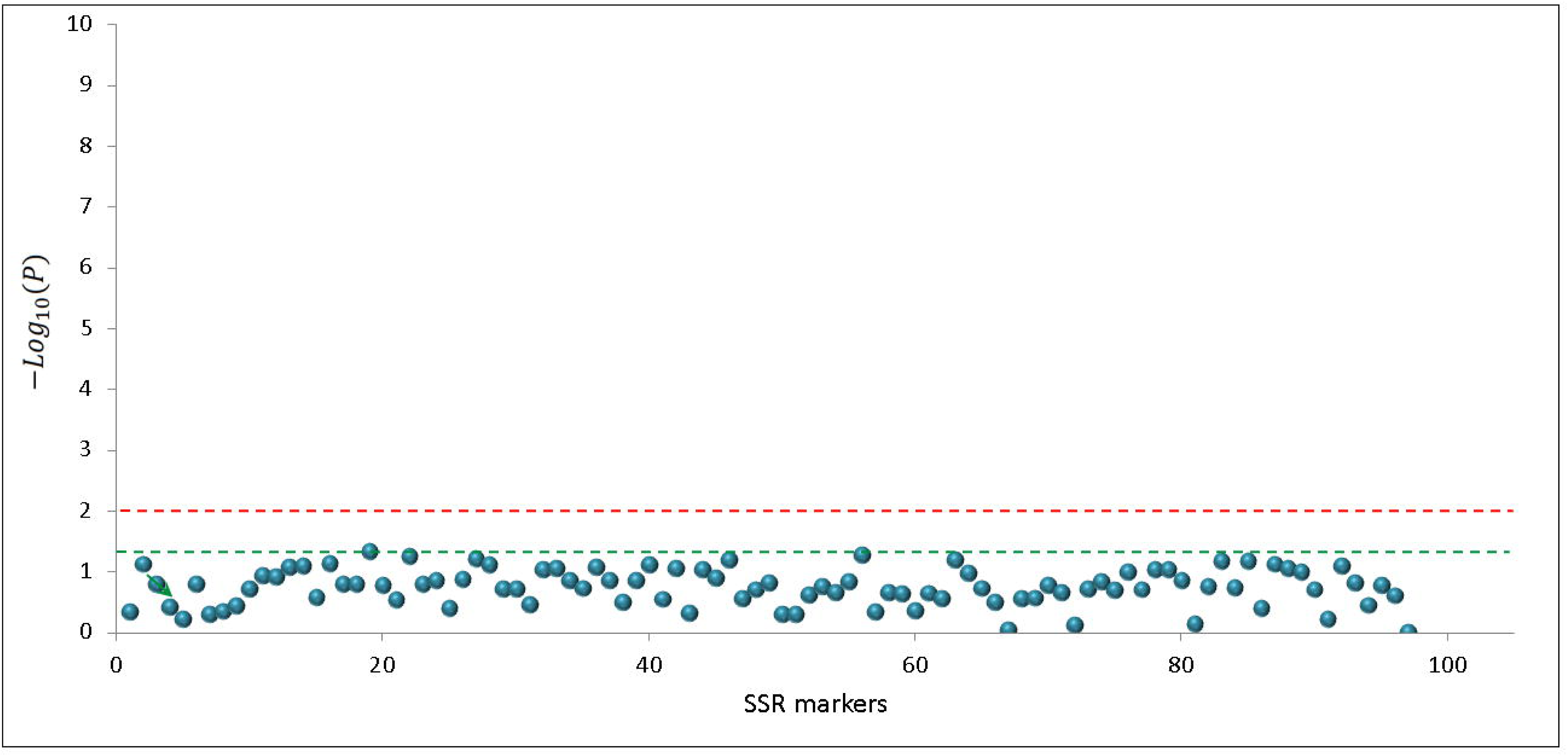
**(a)** Results of genome-wide association scan for leaf shape in diverse set of 406 cotton inbred lines. Blue circles represent the *P*-value of each marker tested in a logistic regression model that also included the first three principal components of the population structure analysis. Three size-based markers from candidate genes are indicated by red arrows while two SNP markers are indicated by green arrows. The green and red dashed lines represent the 0.05 and 0.01 significance levels. **(b)** Results of second genome-wide association scan for leaf shape in diverse set of 406 cotton inbred lines, adjusting for the effects of the candidate gene. Blue circles represent the *P*-value of each marker tested in a logistic regression model that also included the most significant candidate gene marker from the initial scan plus the first three principal components of the population structure analysis. The green and red dashed lines represent the 0.05 and 0.01 significance levels.

**Supplementary Figure 3.**
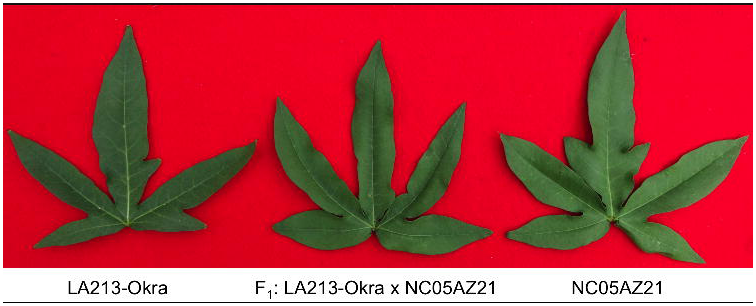
Allelism between *okra* leaf shapes of parental accession (NC05AZ21) and isoline (LA213-*okra*). *Okra* leaf shape genes in isoline LA213-*okra* and NC05AZ21 are allelic. Leaf shape phenotypes of greenhouse grown parents and their F_1_ hybrid at approximately 40 days after planting.

**Supplementary Figure 4.**
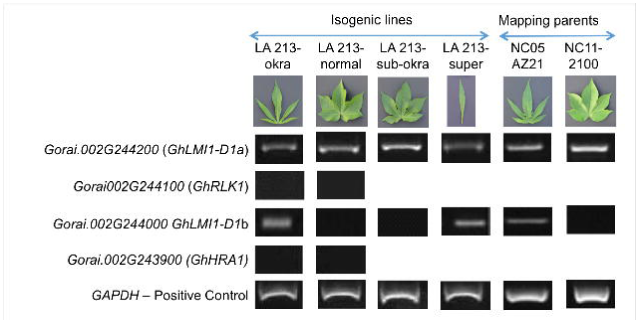
Semi-quantitative expression analysis of leaf shape candidate genes. Neither *GhHRA1* (*Gorai.002G243900*) nor *GhRLK1* (*Gorai.002G244100*) were expressed in critical young leaf tissue, eliminating these two genes from consideration. *GhLMI1-D1a* (*Gorai.002G244200*) appeared equally expressed across leaf shapes. *GhLMI1-D1b* (*Gorai.002G244000*) was expressed only in leaf shapes (*okra* and *super-okra*) with the larger promoter indicating differential expression of this gene could play a major role in leaf shape. Glyceraldehyde-3-phosphate dehydrogenase (*GAPDH*) was the reference gene.

**Supplementary Figure 5.**
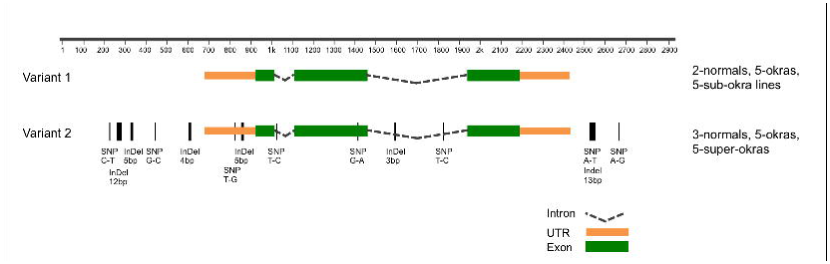
Nucleotide polymorphisms of the *GhLMI1-D1a* gene among different leaf shapes. Two variants of *GhLMI1-D1a* were found, neither of which perfectly associated with leaf shape. Both variants encode full-length proteins and have no obvious polymorphisms that would alter function.

**Supplementary Figure 6.**
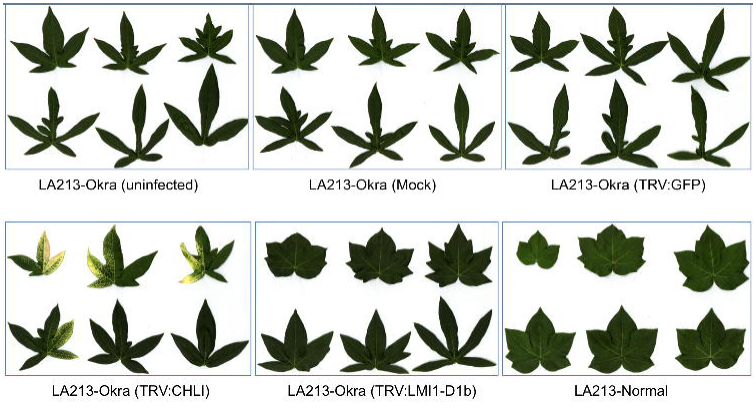
Leaves from representative VIGS plants. True leaves five through ten from representative plants of all VIGS treatments are shown. Severe reductions in leaf lobing and sinus depth was seen in the LA213- *okra* TRV: *LMI1-D1b* that briefly produced normal leaves. Abolishment of viral silencing proceeded similar to that seen in the TRV: CHLI positive control treatment.

**Supplementary Figure 7.**
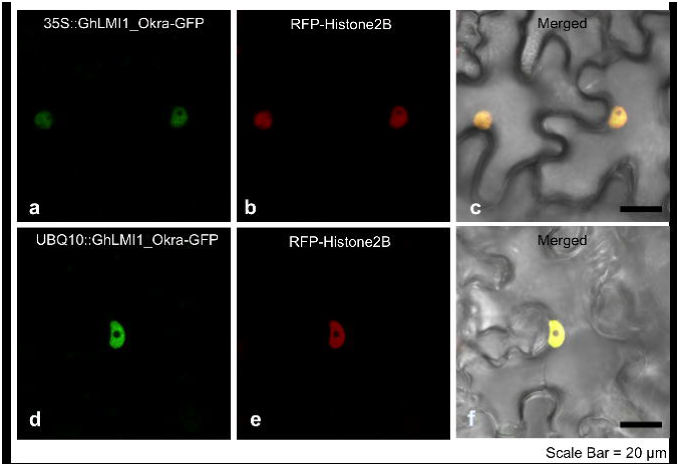
*GhLMI1-D1b^Okra^* localizes to the nucleus. A *GhLMI1-D1b^Okra-^GFP* fusion construct driven by the 35S (**a,b,c**) or pUBQ10 (**d,e,f**) promoter was transiently expressed in *Nicotiana benthamiana* leaves. Signal from *GhLMI1-D1b-^Okra-^GFP* (**a,d**), the nuclear marker *RFP-Histone2B* (**b,e**) or the merged images including brightfield (**c,f**) are shown. Scale bar: 20µm

